# Magnitude-sensitive reaction times reveal non-linear time costs in multi-alternative decision-making

**DOI:** 10.1101/2021.05.05.442775

**Authors:** James A. R. Marshall, Andreagiovanni Reina, Célia Hay, Audrey Dussutour, Angelo Pirrone

## Abstract

Optimality analysis of value-based decisions in binary and multi-alternative choice settings predicts that reaction times should be sensitive only to differences in stimulus magnitudes, but not to overall absolute stimulus magnitude. Yet experimental work in the binary case has shown magnitude sensitive reaction times, and theory shows that this can be explained by switching from linear to geometric time costs, but also by nonlinear subjective utility. Thus disentangling explanations for observed magnitude sensitive reaction times is difficult. Here for the first time we extend the theoretical analysis of geometric time-discounting to ternary choices, and present novel experimental evidence for magnitude-sensitivity in such decisions, in both humans and slime moulds. We consider the optimal policies for all possible combinations of linear and geometric time costs, and linear and nonlinear utility; interestingly, geometric discounting emerges as the predominant explanation for magnitude sensitivity.

## Introduction

While the normative, optimal policy, approach to understanding decision-making is now well established for perceptual decisions (*e*.*g*. ***Bogacz et al. (2006***)), it has only recently been applied to value-based decisions (***Fudenberg et al., 2018***; ***Tajima et al., 2016, 2019***); such decisions differ from perceptual decisions because decision makers are rewarded by the value of the selected option, rather than whether or not they selected the best option (*e*.*g*. ***Pirrone et al. (2014***); ***Tajima et al. (2016***, 2019); ***Krajbich et al. (2010***)). Recently researchers have analysed multi-alternative value-based decision-making (***Tajima et al., 2019***), building on earlier work in optimal decision policies for binary value-based choices (***Tajima et al., 2016***). Through sophisticated analysis based on the standard tool for solving such decision problems, stochastic dynamic programming (***Mangel and Clark, 1988***; ***Houston and McNamara, 1999***), the authors also present neurally-plausible decision mechanisms that may implement or approximate the optimal decision policies (***Tajima et al., 2016, 2019***). These policies turn out to be described by rather simple and well-known decision mechanisms, such as drift-diffusion models with decision thresholds that collapse over time for the binary case (***Tajima et al., 2016***), and nonlinear time-varying thresholds that interpolate between best-vs-average and best-vs-next in the multi-alternative case (***Tajima et al., 2019***).

Interestingly, the theoretically optimal policy for the binary decision case (***Tajima et al., 2016***) is inconsistent with empirical observations of magnitude-sensitive reaction-times (***Teodorescu et al. (2016***); ***Pirrone et al. (2018a***); ***Steverson et al. (2019***); ***Zajkowski et al. (2019***); ***Turner et al. (2019***), but see ***Bhui (2019***)), unless assumptions are made that subjective utilities for decision-makers are nonlinear, or decisions are embedded in a fixed-length time period with known or learnable distributions of trial option values, so that a variable opportunity cost arises from decision time (***Tajima et al., 2016***). Furthermore, even single-trial dynamics lead to magnitude sensitive reaction times (***Pirrone et al., 2018b***).

Previous analyses made an assumption that appears widespread in psychology and neuroscience, that decision-makers should optimise their Bayes Risk from such decisions (***Tajima et al., 2016, 2019***); this is equivalent to maximising the expected value of decisions in which there is a linear cost for the time spent deciding (***Bogacz et al., 2006***; ***Pirrone et al., 2014***). For a lab subject in a pre-defined and known experimental design this may appear appropriate, for example because there may be a fixed time duration within which a number of decision trials will occur and the subject can learn the value distribution of the trials (e.g. ***Bogacz et al. (2006***); ***Pirrone et al. (2014***)). However, an alternative and standard formulation of the Bellman equation, the central equation in constructing a dynamic program, accounts for the cost of time by discounting future rewards geometrically, so a reward one time step in the future is discounted by rate *γ* < 1, two time steps in the future by *γ*^2^, and so on (see Materials and Methods). This is a standard assumption in behavioural ecology (***Mangel and Clark, 1988***; ***Houston and McNamara, 1999***), in which discounting the future means that future rewards are not guaranteed but are uncertain, due to factors such as interruption, consumption of a food item by a competitor, mortality, and so on. Thus discounting the future represents the inherent uncertainty that animals must make decisions under in their natural environments, in which their brains evolved. The appropriate discount is then the probability that future rewards are realised, hence geometric discounting is optimal since probabilities multiply. Indeed there is extensive evidence of such reward discounting in humans and other animals (*e*.*g*. ***Sellitto et al. (2010***), although this frequently suggests hyperbolic rather than geometric discounting, a fact that in itself merits an explanation based on optimality theory (***McNamara and Houston, 2009***)).

Rederiving optimal policies to account for geometric (***Marshall, 2019***) or general multiplicative (***Steverson et al., 2019***) costs of time qualitatively changes them in the binary decision case, introducing magnitude-sensitive reaction times (***Marshall, 2019***; ***Steverson et al., 2019***). However, disentangling these from nonlinear subjective utility is challenging, and cannot be excluded as an explanation for previous results (***Teodorescu et al., 2016***; ***Pirrone et al., 2018a***; ***Steverson et al., 2019***; ***Zajkowski et al., 2019***; ***Turner et al., 2019***; ***Bhui, 2019***; ***Smith and Krajbich, 2019***; ***Pirrone and Gobet, 2021***).

Here for the first time we extend the theoretical and experimental study of magnitude-sensitivity to three-alternative decisions. We first present evidence for magnitude-sensitive reaction times in three-way equal-alternative decisions. We then present optimal policy analyses and novel numerical simulations for such ternary decisions, both in human subjects undertaking a psychophysical task, and unicellular organisms engaged in foraging. Importantly, for a wide variety of utility functions, strong magnitude-sensitivity is only observed when there is a multiplicative cost for time, rather than the previously assumed linear time cost. Thus magnitude-sensitivity is revealed as genuinely diagnostic for multiplicative time costing, as all other assumptions either do not generate this phenomenon, or can be discounted.

### Results

As we were testing theory developed to explain decision-making by animals with brains, we conducted psychophysical experiments with human subjects. However, we also conducted foraging experiments with a unicellular slime mould; testing theory across multiple species and behavioural tasks increases confidence when multiple agreements with theory are observed (***Pirrone et al., 2018a***), and slime moulds have become a model system, with multiple experiments seeking to reproduce behavioural predictions from neuroscience and psychology (***Latty and Beekman, 2011***; ***Reid et al., 2016***; ***Dussutour et al., 2019***).

Multi-Alternative Decisions in Human Psychophysical Trials are Magnitude-Sensitive Here we provide strong empirical evidence for magnitude sensitivity with multiple alternatives in humans, using an experimental paradigm similar to the one used to show magnitude sensitivity for two-alternative decision making (***Teodorescu et al., 2016***; ***Pirrone et al., 2018a***). Details of the experiment (methods, participants, etc.) are reported in Materials and Methods. Participants had to choose which of three above-threshold grey patches was brighter in an online experiment. Although the experiment included conditions for which a brighter alternative existed, conditions of interest were equal alternatives of different magnitude, that is, conditions for which the three patches had the same brightness that could vary across magnitude conditions. Equal alternatives allow us to test hypotheses regarding magnitude sensitivity, by keeping differences in evidence constant (***Pirrone et al., 2018a***,b; ***Pirrone and Gobet, 2021***).

As previously done for binary decisions (***Pirrone et al., 2018a***,b), here we focused our analyses exclusively on equal alternatives. For the analyses, we did not censor any datapoints.

As shown in Figure 1, the data show strong magnitude sensitivity, given that choices for equal alternatives of higher magnitude conditions (higher brightness on a scale from 0 to 1 on Python) were made faster.

**Figure 1.**
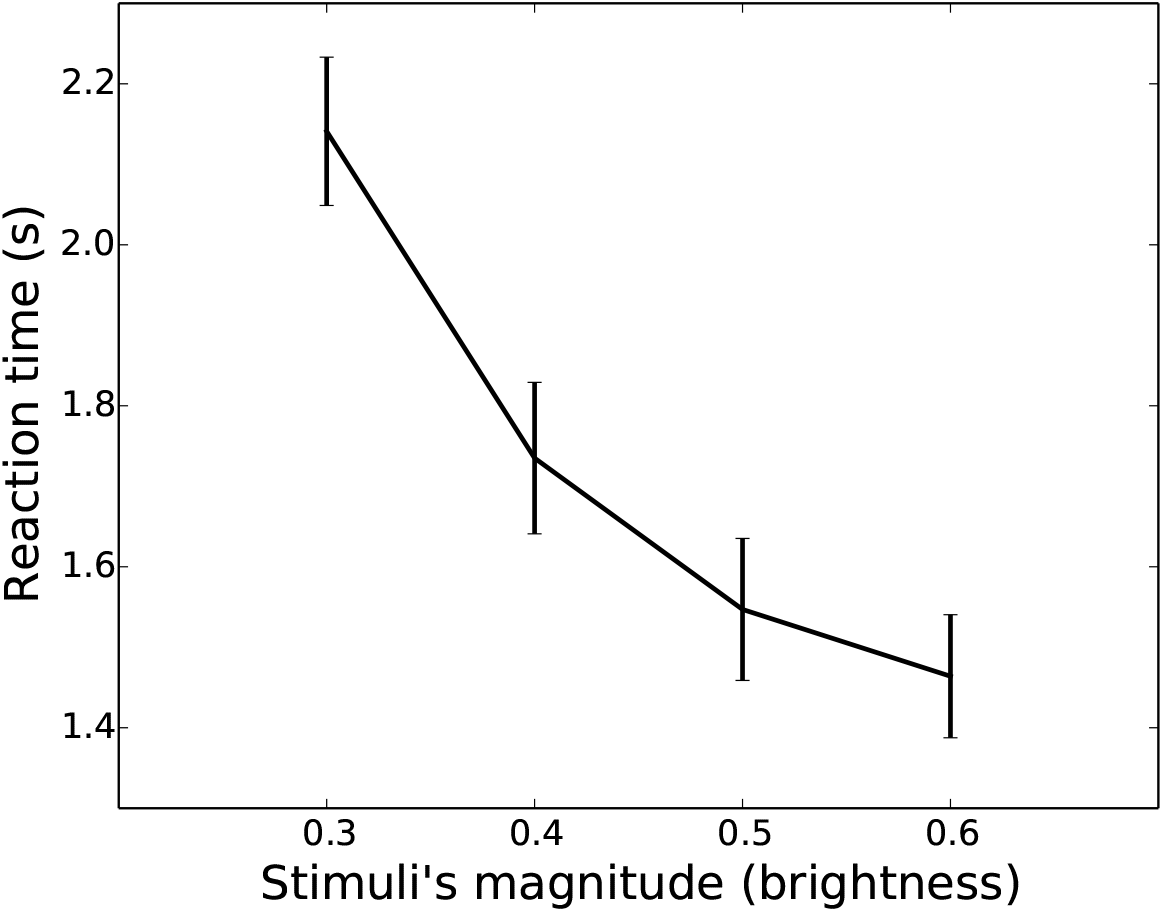
Empirical results from the behavioural online experiment. Decreasing reaction times as a function of the magnitude of the equal alternatives. X-axis presents mean brightness of equal alternatives (0.3, 0.4, 0.5, 0.6), on a scale of brightness from 0 to 1 in PsychoPy. Y-axis presents mean reaction times, in seconds. Bars show 95% confidence intervals. Participants experienced equal alternative conditions, interleaved with unequal alternative trials in pseudo-randomised order. Participants that performed the whole experiment experienced each equal alternative presentation ten times.

To assess if reaction times decreased as a function of the mean brightness of the equal alternatives, we used a linear mixed model in R. The model was fitted by specifying as fixed effect (explanatory variable) the brightness of equal alternatives as a continuous predictor. The participant ID was also added to the model as a factor for random effects. Reaction times significantly decreased as a function of the mean brightness of the alternatives (b = -1.95, p < .001, CI -2.14 -1.75). Further details for the mixed-effect regression are presented in the supplementary information (Table S1).

As the COVID-19 pandemic necessitated an online experiment we could not collect or control information on a number of possible confounds (viewer position, motivation, room luminosity, etc.), and there are multiple sources of unaccounted variability in our online experiment; however there is no *a priori* reason to expect these to act as consistent confounds in the magnitude-sensitive reaction times observed. Furthermore, the very large sample size for our study (*N* = 117; compared to *N* = 8 and *N* = 9 for previous similar investigations (***Teodorescu et al., 2016***; ***Pirrone et al., 2018a***)) should minimise effects due to randomly-distributed confounds.

#### Multi-Alternative Decisions in Foraging Trials by Unicellular Organisms are Magnitude-Sensitive

Here, using slime moulds of the species *Physarum polycephalum*, we also observed strong empirical evidence for magnitude sensitivity with three alternative foraging tasks. Details of the experiment are reported in Materials and Methods. Slime moulds were confronted with a choice offering three equal food sources. We increased the magnitude of the options by increasing the quality of the food sources. As shown in Figure 2, the latency to reach one of the alternatives depends on the quality of the food sources; the higher the quality the faster the slime mould. This was confirmed by a linear mixed model similar to the one applied to the human data, in which reaction times significantly decreased as a function of food quality (b = -0.03, p < .001, CI -0.03 -0.02; further details, Table S2 in the supplementary information).

**Figure 2.**
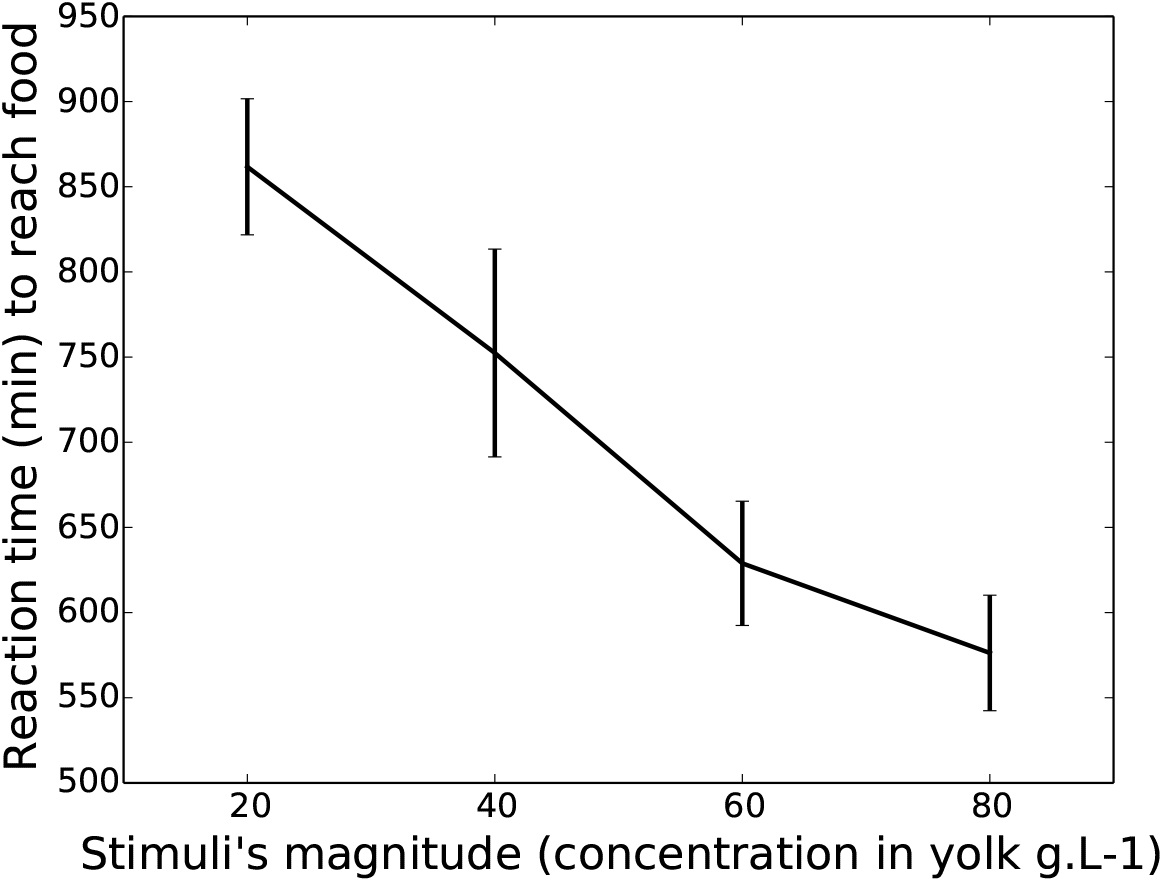
Empirical results from the slime mould experiment. Decreasing latencies to reach a food source as a function of the magnitude of the equal alternatives. X-axis presents the concentration in egg yolk of equal food sources (20, 40, 60, 80 g.L-1). Y-axis presents mean latency to reach a food source, in minutes. Bars show 95% confidence intervals. 50 slime moulds were tested for each magnitude for a total of 200 slime moulds.

### Optimal Policies

For our theoretical analysis we begin by re-deriving optimal policies for decisions when the change is made from linear costing of time, or Bayes Risk, to geometric discounting of future reward. Note that geometric discounting of future rewards is similar to, but not the same as, non-linear utility. As remarked in the introduction above, for binary decisions magnitude-sensitive reaction times can be explained by optimal decision policies for either multiplicative (*e*.*g*. geometric) time discounting (***Marshall, 2019***; ***Steverson et al., 2019***) or nonlinear subjective utility with linear time costs (***Tajima et al., 2016***). In the multi-alternative case, on the other hand, the picture is more nuanced; moving from linear costing of time to geometric discounting of future rewards changes complicated time-dependent non-linear decision thresholds ((***Tajima et al., 2019***) Fig. 7) into either simple linear ones that collapse over time for lower-value option sets (Fig. 3), or nonlinear boundaries that evolve over time similarly to the Bayes Risk-optimising case for higher-value option sets (***Marshall (2019***); Fig. 3). As Tajima *et al*. note, the simpler linear decision boundaries implement the ‘best-vs-average’ decision strategy, whereas the more complex boundaries interpolate between ‘best-vs-average’ and ‘best-vs-next’ decision strategies (***Tajima et al., 2019***); interestingly simply moving to nonlinear subjective utility with linear time costs simplifies the decision strategy to the ‘best-vs-next’ strategy (Fig. 3; see ***Tajima et al. (2019***), Fig. 6C).

**Figure 3.**
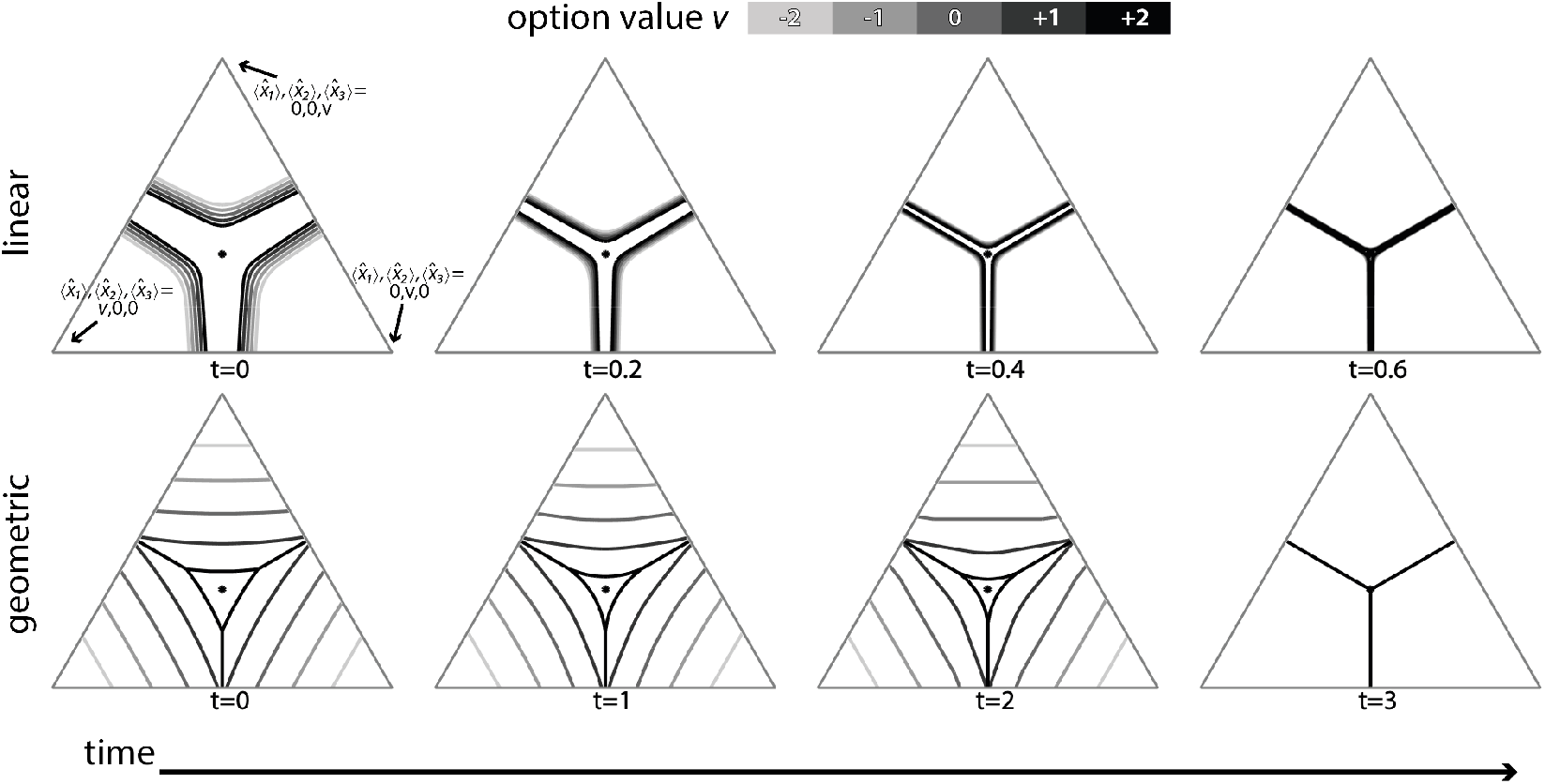
Linear time costs lead to weakly magnitude-sensitive optimal policies (top row), while geometric discounting of reward leads to strongly magnitude-sensitive optimal policies (bottom row). In the linear time cost (Bayes Risk) case nonlinear subjective utility changes complex time and value-dependent decision boundaries in estimate space into a simple mostly magnitude-insensitive ‘best-vs-next’ strategy (top row; see ***Tajima et al. (2019***), Fig. 6C). For geometric discounting of rewards over time, optimal decision boundaries are strongly magnitude-sensitive and interpolate between simple ‘best-vs-average’ and ‘best-vs-next’ strategies (see ***Tajima et al. (2019***), Fig. 6). Triangles are low dimensional projections of the 3-dimensional evidence estimate space onto a plane moving along the equal value line, at value *v* (***Tajima et al., 2019***). Dynamic programming parameters were: prior mean 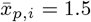 and variance 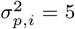, waiting time *t*_*w*_ = 1, temporal costs *c* = 0, *γ* = 0.2, and utility function parameters *m* = 4, *s* = 0.25 (for the linear time cost) and *m* = 4, *s* = 3.5 (for the geometric time cost).

#### Multi-Alternative Decisions: Optimal Policies are Weakly Magnitude-Sensitive for Nonlinear Subjective Utility Under Bayes Risk-Optimisation

Under Bayes Risk-optimisation it is known that, for binary decisions, optimal policies are magnitude-insensitive when subjective utility is linear, whereas they are magnitude-sensitive when subjective utility is nonlinear (***Tajima et al., 2016, 2019***).

For ternary decisions, however, even with nonlinear subjective utility, policies exhibit very weak magnitude-sensitivity early in decisions, becoming magnitude-insensitive as decisions progress (Fig. 3, row ‘linear‘). Sensitivity analysis shows that magnitude-insensitivity is a general pattern. An informal understanding of this can be arrived at by appreciating that sigmoidal functions have two extremes of parameterisation; in one extreme they are almost linear, hence will be mostly magnitude insensitive due to the known result (***Tajima et al., 2016***). At the other extreme, the function becomes step-like; in this case options are either good or bad, and the optimal policy rapidly becomes ‘choose the best’ (Fig. 4), since under such a scenario sampling is of minimal benefit as early information quickly indicates whether an option is good or bad, and choosing the first option that appears to be good is optimal.

**Figure 4.**
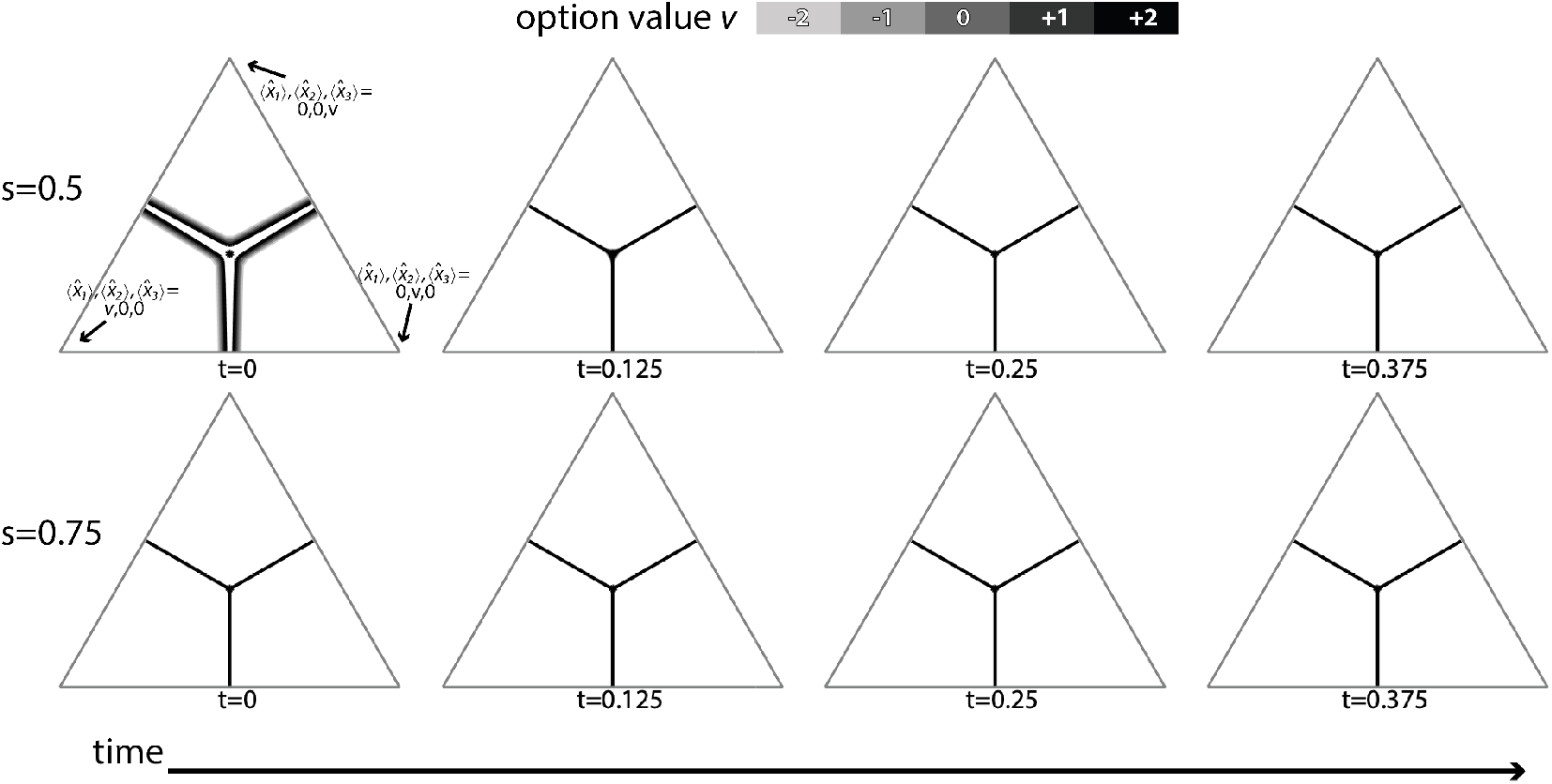
Optimal policies for linear time cost (Bayes Risk) rapidly transition from approximately linear subjective utility, and hence weakly magnitude-sensitive, decision boundaries in estimate space (Fig. 3, top row for *s* = 0.25; present figure, top row for *s* = 0.5, to more step-like subjective utility where immediate ‘choose the best’ decision-boundaries are necessarily magnitude-insensitive (bottom row for *s* = 0.75, and higher values of *s*). Triangles are low dimensional projections of the 3-dimensional evidence estimate space onto a plane moving along the equal value line, at value *v* (***Tajima et al., 2019***). Dynamic programming parameters were: prior mean 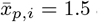 and variance 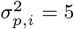, and utility function parameters *m* = 4, *s* ∈ {0.5, 0.75}.

#### Multi-Alternative Decisions: Optimal Policies Become Magnitude-Sensitive Under Geometric Discounting

As previously shown (***Marshall, 2019***; ***Steverson et al., 2019***), assuming geometric temporal discounting, the optimal policy for binary decisions is magnitude-sensitive. In ternary decisions, geometric discounting has the same effect; regardless of utility function linearity, the optimal policy is magnitude-sensitive (Fig. 3, row ‘geometric‘).

### Numerical Simulations

Since noise-processing is fundamental in determining reaction times, we confirmed the results on magnitude sensitivity from our optimal policy analysis via numerical simulation of Bayes-optimal evidence-accumulating agents using those policies (see Materials and Methods). These numerical simulations confirmed the qualitative results from the optimal policy analysis; reaction times for ternary decisions under linear time costing are only weakly magnitude sensitive even for nonlinear subjective utility functions, while under geometric time costing reaction times become strongly magnitude sensitive for most utility functions examined.

#### Multi-Alternative Decisions: Simulated Reaction Times are Weakly Magnitude-Sensitive for Nonlinear Subjective Utility Under Bayes Risk-Optimisation

Across all nonlinear subjective utility functions considered, linear time costing resulted in weakly magnitude-sensitive simulated reaction times (Fig. 5). This agrees with the weak magnitude-sensitivity observed in the optimal policies derived above (Fig. 3). Note, however, that this contrasts with the binary decision case in which optimal policies, and hence reaction times, become magnitude sensitive under linear time cost when subjective utility is nonlinear (***Tajima et al., 2016***). An informal justification for this is given above in analysis the optimal decision boundaries computed via dynamic programming.

**Figure 5.**
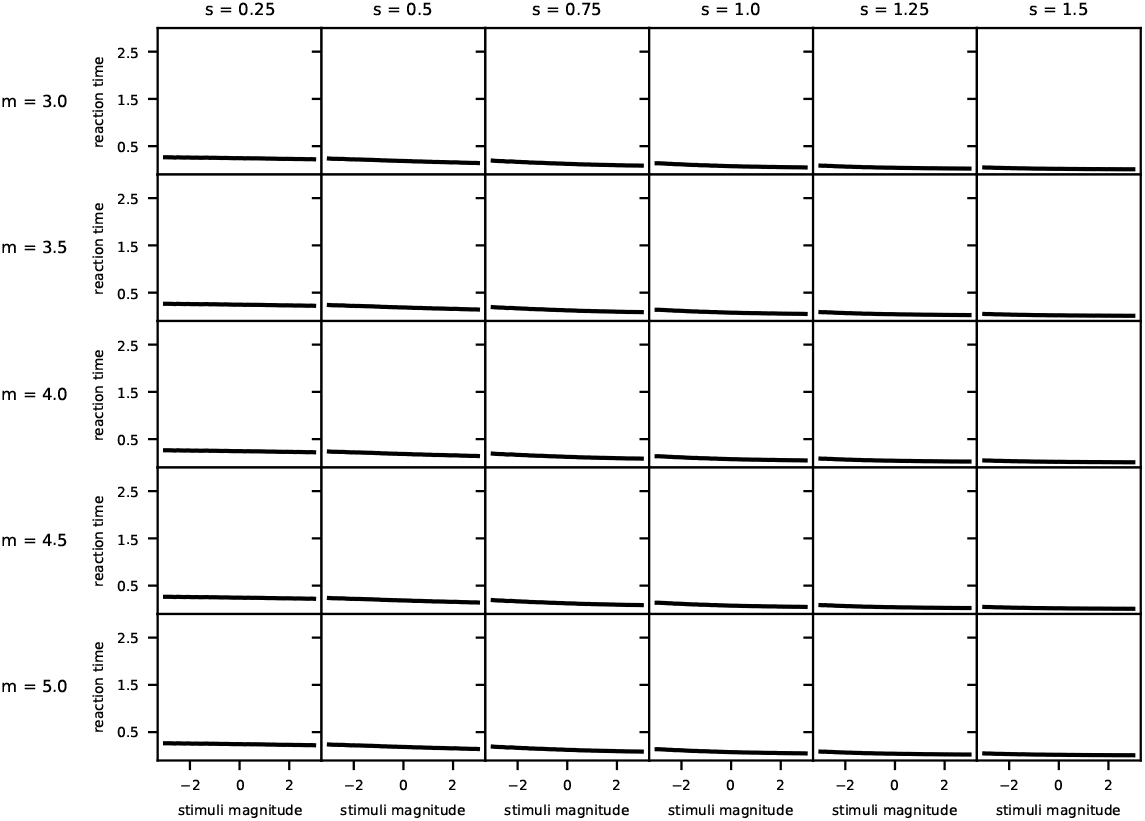
Linear time costs lead to weakly magnitude-sensitive simulated reaction times across a range of nonlinear subjective utility functions for equal value option sets. Simulation parameters were: prior mean 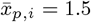 and variance 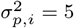, observation noise variance 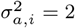, temporal cost *c* = 0, waiting time *t*_*w*_ = 1, and simulation timestep *dt* = 5 × 10^−3^. Lines are the mean reaction time for 10^4^ simulations, 95% confidence intervals are shown as red shading (mostly invisible because smaller than the linewidth).

#### Multi-Alternative Decisions: Simulated Reaction Times Become Magnitude-Sensitive Under Geometric Discounting

In contrast to linear time costing, across all nonlinear subjective utility functions considered, geometric time costing resulted in strongly magnitude sensitive simulated reaction times (Fig. 6), with longer reaction times for lower value equal-value option sets; this strategy was previously hypothesised to be optimal (***Pais et al., 2013***). The strong magnitude-sensitivity in the numerical simulations corresponds with the strong magnitude-sensitivity observed in the optimal policies derived above (Fig. 3).

**Figure 6.**
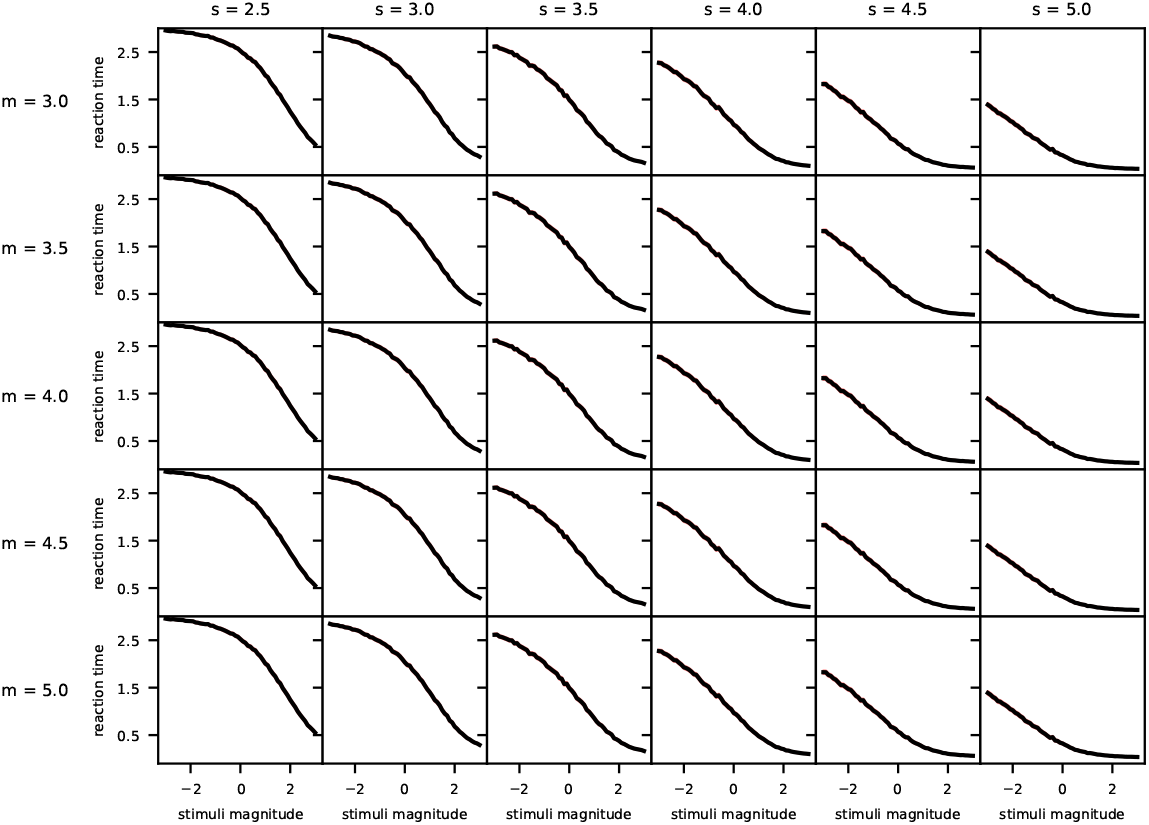
Geometric discounting of reward leads to strongly magnitude-sensitive simulated reaction times across a range of nonlinear subjective utility functions, with decisions postponed for low equal-value option sets. Simulation parameters were: prior mean 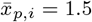 and variance 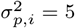, observation noise variance 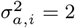, temporal cost *γ* = 0.1, and simulation timestep *dt* = 5 × 10^−3^. Lines are the mean reaction time for 10^4^ simulations, 95% confidence intervals are shown as red shading (mostly invisible because smaller than the linewidth).

## Discussion

In understanding behaviour, which is a product of evolution, searching for optimal algorithms for typical decision problems can provide great insight. This normative approach can explain observed behaviours, and predict new behavioural patterns, based on evolutionary advantage. Yet the assumptions underlying such model analyses can prove crucial. Recently it has been asked what optimal decision algorithms look like for multi-alternative value-based choices, in which subjects are rewarded not by whether their decision was correct or not, but by the value to them of the selected option (***Tajima et al., 2019***). The resulting algorithms correspond to earlier simple models for perceptual and value-based decision-making. These findings, however, rest on an assumption that time is a linear cost for subjects. Here we have shown that deciding human subjects and foraging unicellular organisms do, however, exhibit marked magnitude sensitivity in ternary decisions, as previously shown for binary decisions (***Pirrone et al., 2018a***; ***Dussutour et al., 2019***). We have also shown that optimality theory that discounts future rewards multiplicatively based on time is the foremost explanation for such observations of magnitude-sensitivity; nonlinear subjective utility alone is not sufficient to give rise to strongly magnitude-sensitive decision times when time is treated as a linear cost.

### Behavioural Predictions

The Bayes Risk optimal policy is approximated by a neural model that is consistent with observations of economic irrationality (***Tajima et al., 2019***), hence it will be important to see if a revised neural model based on the revised optimal policy still shows such agreement. For example, while in the binary case magnitude-sensitive reaction times can be explained both by nonlinear subjective utility functions, and by multiplicative discounting rather than Bayes Risk, in the multi-alternative case our analysis suggests that the same phenomenon is explained *primarily* by multiplicative discounting of future rewards and not by nonlinear utility.

### Optimality Criteria

Practitioners of behavioural ecology have established principles to deal with empirically-observed deviations from the predictions of optimality theory (***Parker and Smith, 1990***); two of the most useful are to consider that the optimisation criterion has been misidentified, or the behaviour in question is not really adaptive. Tajima and colleagues employ an exemplary approach, attempting to combine the best of the approaches of normative and mechanistic modelling (***McNamara and Houston, 2009***); yet it bears remembering that subjects may not be trying optimally to solve the simple decision problem they are presented in the lab, but rather making use of mechanisms that evolved to solve the problem of living in their natural environment (***Fawcett et al., 2014***); indeed the experimental data presented here were produced by subjects who received no reward, yet nevertheless acted as if they were making an economic, value-based, decision rather than a purely perceptual, accuracy-based one.

## Materials and Methods

### Psychophysical Experiment

#### Participants

This experiment was conducted during the COVID-19 pandemic, from May 13th – June 1st 2020.. Given that it was not possible to recruit participants for a laboratory experiment, we instead recruited them online using Pavlovia (***Peirce et al., 2019***; ***Peirce, 2007***), an online platform for psychophysical experiments implemented in PsychoPy.

Running a perceptual experiment online has a number of limitations: first, there is no way to ensure that participants are focused on the task and minimising distractions - to mitigate this we kept the task short and participants were instructed to concentrate on it; second, Pavlovia (as of March-May 2020) only allows stimuli to be drawn in units relative to window size (i.e., the window in which the experiment is displayed) or in pixels, hence their size and position relative to the fixation cross will vary across devices, depending on specific window sizes. However, even if the size and location of stimuli could be kept constant across participants, participants’ distance from the screen cannot be controlled during an online experiment.

While in previous two-alternative experiments (***Teodorescu et al., 2016***; ***Pirrone et al., 2018a***) a limited number of participants (N < 10) performed a large number of trials, for our online experiment we aimed at a large number of participants performing a limited number of trials. This strategy is beneficial for online studies since the large number of participants helps ensure that variation in participants’ motivation or viewing arrangements is averaged out. We therefore recruited 117 participants via external advertisement on Twitter, and internal email lists at the University of Sheffield (mean age = 40.4, SD = 11.2774, range 23 -77; 79 females, 37 males, 1 did not indicate their gender). We requested participants to follow the link to the experiment only if aged 18 years or older. The experiment lasted about 5 minutes and participation was voluntary; participants did not receive any reward for their participation.

After reading the instructions, participants were informed that by continuing they were confirming that they understood the nature of the experiment and consented to participate. Participants were also informed that they could leave the experiment at any time by closing the browser. For this experiment, all procedures were approved by the University of Sheffield, Department of Computer Science Ethics Committee.

#### Experimental setup

Similarly to previous studies (***Teodorescu et al., 2016***; ***Pirrone et al., 2018a***), stimuli consisted of three homogeneous, round, white patches in a triangular arrangement on a grey background, as depicted in Figure 7. Throughout the task participants were presented with a central fixation cross that they were requested to fixate on.

**Figure 7.**
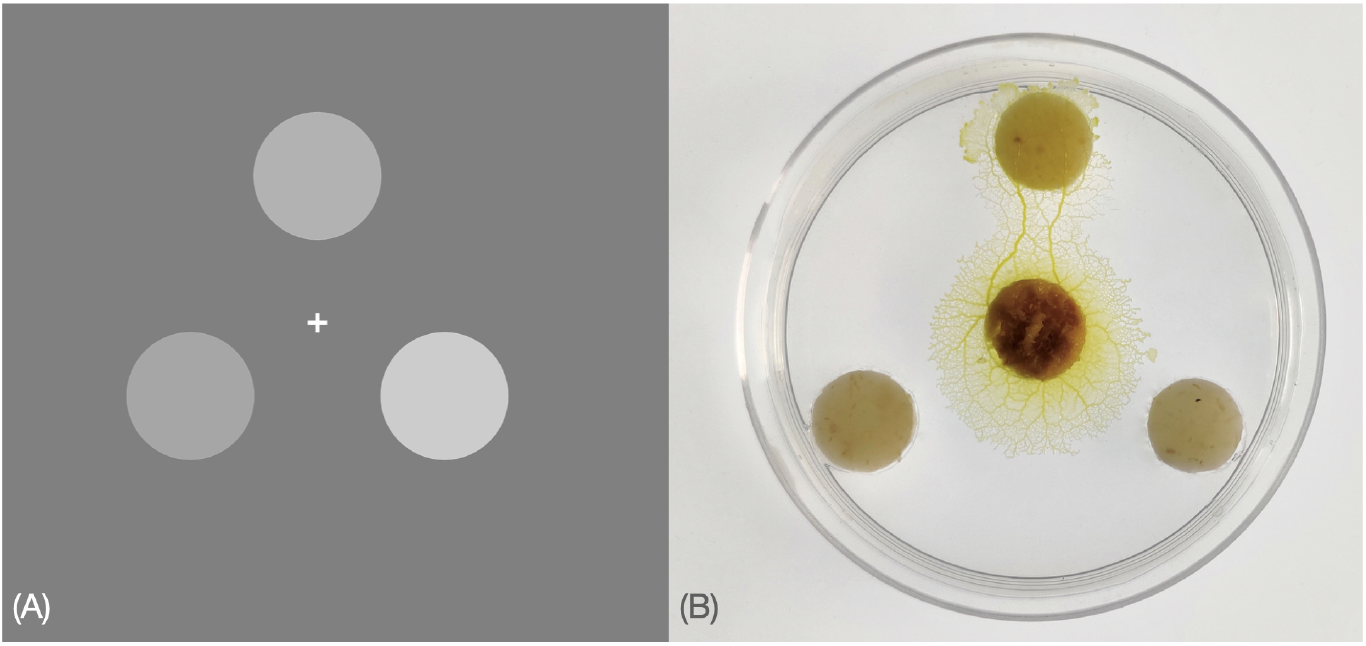
(A) Stimuli example for human psychophysical experiments: Participants were requested to decide as fast and accurately as possible which of the three stimuli was brighter; they were asked to maintain fixation on the cross at the centre of the screen and minimise distraction for the short duration of the experiment. Unknown to participants, conditions of interest were conditions for which the stimuli had equal mean brightness. (B) Photograph showing a slime mould that chose one food alternative among three equal ones. The slime mould was placed in the centre of a petri dish (6cm ø) filled with agar gel (10 g/l) at a distance of 2cm from each food alternative.

On a scale from 0 to 1 in PsychoPy, the patches could have a brightness of 0.3, 0.4, 0.5 or 0.6. There were 4^3^ = 64 possible trial combinations, of which 4 were equal alternatives (i.e., alternatives having a brightness of [0.3,0.3,0.3], [0.4,0.4,0.4], [0.5,0.5,0.5] or [0.6,0.6,0.6]). We selected 10 equal trial repeats, and only one trial repeat for all possible unequal alternatives, for a total of 100 trials per participant. On each frame, a Gaussian random variable with mean 0 and standard deviation of 0.25 × (mean brightness of the alternative) was added separately to the brightness level of each patch; the signal-to-noise ratio was thus kept constant across equal alternatives of different magnitude. The order of presentation of the alternatives was pseudo-randomised across participants and there was no systematic link between patch position and best option.

Three grey patches were presented simultaneously on the screen and subjects were asked to decide which of the three was brighter by pressing ‘left’, ‘right’ or ‘up’ on a keyboard using their second, fourth or third right-hand fingers, via a line-drawn diagram of a hand over a keyboard presented before the experiment began; specific instructions for left-handed participants were not provided, and we did not record handedness. The inter-trial interval, during which participants were presented with only the fixation cross, was selected at random between 0.5 seconds, 1 second or 1.5 seconds for each trial. Subjects were instructed to be as fast and accurate as possible and to maintain their fixation on the cross at the centre of the screen throughout the experiment. Before the experiment they were presented with 6 training trials (unequal alternatives) to familiarise themselves with the task. Participants were not provided with any feedback after each trial and were not informed about the presence of the equal-alternatives conditions.

### Slime Mould Experiment

*Physarum polycephalum*, also known as the acellular slime mould, is a giant polynucleated single cell organism that inhabits shady, cool, and moist areas. In the wild, *P. polycephalum* eats bacteria and dead organic matter. In the presence of chemical stimuli in the environment, slime moulds show directional movements (*i*.*e*. chemotaxis).

Slime moulds of strain LU352 kindly provided by Professor Dr Wolfgang Marwan (Max Planck Institute for Dynamics of Complex Technical Systems, Magdeburg, Germany) were used for the experiments. Slime moulds were initiated with a total of 10 sclerotia which are encysted resting stages. The sclerotia were soaked in water and placed in petri dishes (140 mm ø) on agar gel (1%). Once revived, slime moulds start to explore the agar gel, usually 24h after the reactivation of the sclerotia. The slime moulds were then reared for a month on a 1% agar medium with rolled oat flakes (Quaker Oats Company®) in Petri dishes (140 mm ø). They were kept in the dark in a thermoregulated chamber at a temperature of 20 degrees Celsius and a humidity of 80%. The day before the experiment the slime moulds were transferred on a 10% oat medium (powdered oat in a 1% agar solution) in Petri dishes (diameter 140 mm). The experiments were carried out in a thermoregulated chamber and pictures were taken with a Canon 70D digital camera.

Slime moulds were presented with a choice between three equal food sources in an arena consisting of 60 mm diameter Petri dish filled with plain 1% agar. We punched three holes (10mm ø) in the arena and filled them with a food source (10mm ø). We used four different food patches varying in quality: 2% w/v powdered oat mixed with either 2, 4, 6 or 8% w/v egg yolk. Once the food sources were set in each hole, we placed a slime mould (10mm ø) in the centre of the arena 2cm away from each food. We replicated the experiment 50 times for each food quality. For each replicate, we measured the time taken by the slime mould to reach either one of the three food sources.

To assess the difference in the latency to reach the food as a function of the food quality, we used a linear mixed model (function lmer, Package lme4) in R (RStudio Version 1.2.1335). The models were fitted by specifying the fixed effects (explanatory variables) the concentration in yolk (continuous predictor). The sclerotia identity was also added to the model as a random factor. We transformed the dependant variable using the “bestNormalize” function (“bestNormalize” package). The outcome of the model is presented in the supplementary information (Table S2).

#### Optimal policies

Optimal policy computations were performed in Matlab (***MathWorks, 2020***), and were adapted from the dynamic program of ***Tajima et al. (2019***). Optimal policies were computed for Bayes Risk and geometric discounting, for linear utility (*r ≔ x*), and for non-linear utility functions having the form

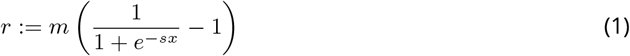

where *m* and *s* are shape parameters for the logistic function determining the interval of utilities and steepness of the slope respectively, and *x* is the raw input value. We systematically varied the *m* and *s* parameters to test magnitude-sensitivity under Bayes Risk optimisation and geometric discounting, under a range of utility function shapes ranging from almost linear, to almost stepwise. Note that a sigmoid curve includes an interval in which subjective utility is an accelerating function of input value when the latter is negative, and an interval in which it is a decelerating function when the latter is positive; thus testing for magnitude-sensitivity over the full interval of raw input values tests a variety of utility function shapes over sub-intervals.

The Bellman equation used in the dynamic programming analysis for the Bayes Risk-optimisation case was

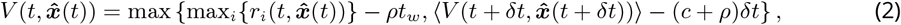

where 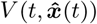 is the value of the state estimates vector 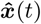 at time *t*, 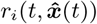 is similarly the expected reward from choosing the *i*-th reward, *δt* is the time interval to the next decision point, *c* is the linear cost per unit time, *ρ* is the reward rate per unit time based on optimal decision-making over a sequence of trials, *t*_*w*_ is the inter-trial waiting time, and ⟨ … ⟩ is expectation over the next time interval (*δt*) (***Tajima et al., 2019***). For the results presented here we set *c* = 0, *t*_*w*_ = 1 and found the optimal *ρ*> 0 using the methods of ***Tajima et al. (2019***); note, however, that since the prior was not varied this reward-rate optimisationcould not induce magnitude-sensitive reaction times in itself.

For the geometric discounting case the Bellman equation becomes

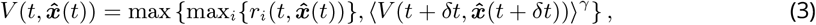

where 0 <*γ* < 1 is a discount factor for rewards received in future timesteps; this discount factor is per-unit-time, hence to discount a reward *δt* < 1 timesteps in the future the appropriate factor is *γ*^*δt/*1^ = *γ* ^*δt*^.

#### Stochastic simulations

Since noise processing is important in determining reaction times, we derived optimal decision policies as above, then tested them through numerical analysis of stochastic models. To test for magnitude-sensitive decision-making we examined the case of *n* = 3 equal-quality alternatives, in which we varied the magnitude of the (equal) stimuli values. Through these stochastic simulations, we tested the impact of the different temporal discount methods—linear or geometric—and of different utility functions—linear or nonlinear—on the decision speed (reaction time RT). The stochastic models simulate the sequential accumulation of evidence for the three alternatives *i* ∈ {1, 2, 3} that are used to compute the expected rewards 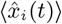. The three evidence estimates can be represented, with a little abuse of notation, as a vector 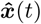 denoting a point in a cube that represents the estimate space. In that cube, we also include the decision boundaries computed as above, indicating the separation of the estimate space into decision regions; in one region continuing to sample is expected to maximise utility, and in the other region taking a decision for the leading option is expected to be the best action.

In our simulations, each time step of length *dt* the decision-maker accumulates three pieces of evidence, one for each option. Evidence for an option *i* is sampled from the normal distribution 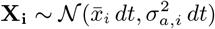, where each sample *x*_*τ,i*_ is a piece of momentary evidence at a small timestep of length *dt* and with sequential index *τ*, 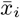 is the true raw value (before any nonlinear utility transformation) of option *i* and 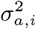 is the variance in accumulation of evidence for *i* (***Tajima et al., 2016***). Before observing any evidence, the decision maker has prior mean and variance, 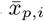 and 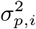 for the distribution of **X**_**i**_. Each new piece of accumulated evidence is used by the decision maker to update the posterior expected reward as

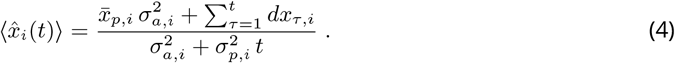

## Code and Data Availability

Optimisation codes for the results presented here can be downloaded from https://github.com/DiODeProject/MultiAlternativeDecisions/tree/stochastic-sim Experimental code and data for the results presented here can be downloaded from https://osf.io/8jumk/

## Acknowledgements

We thank Satohiro Tajima for sharing the code for the binary decision model. Dr Tajima was an exceptionally promising scientist who is sadly missed. We thank Jan Drugowitsch and Alex Pouget for discussions of their results and our own, and Thomas Bose, Nathan Lepora and Sophie Baker for comments on an earlier draft.

