## Supplementary Information for "Magnitude-sensitive reaction times reveal non-linear time costs in multi-alternative decision-making"

| | **Reaction time** | | |
|---|---|---|---|
| *Predictors* | *Estimates* | *CI* | *p* |
| (Intercept) | 0.87 | 0.73 – 1.02 | **<0.001** |
| Brightness | -1.95 | -2.14 – -1.75 | **<0.001** |
| **Random Effects** | | | |
| $\sigma^2$ | 0.58 | | |
| $\tau_{00}$ ID Particiapnt | 0.38 | | |
| ICC | 0.39 | | |
| N ID Participant | 117 | | |
| Observations | 4644 | | |
| Marginal $R^2$ / Conditional $R^2$ | 0.047 / 0.422 | | |

**Table S1** Mixed-effect regression for reaction times as a function of the brightness of the equal alternatives in the human study. Participant ID was included as a random factor. The regression was performed using R (RStudio Version 1.2.1335; function *lmer*, package lme4). Given the typical skewness of reaction times, the dependant variable was transformed (i.e., normalized) using the *bestNormalize* function in R. As the brightness of equal alternatives increased, reaction times significantly decreased.

| | Latency to reach the food | | |
|---|---|---|---|
| *Predictors* | *Estimates* | *CI* | *p* |
| (Intercept) | 1.27 | 0.97 – 1.56 | **<0.001** |
| Food Quality | -0.03 | -0.03 – -0.02 | **<0.001** |
| **Random Effects** | | | |
| $\sigma^2$ | 0.65 | | |
| $\tau_{00}$ ID Plasmodium | 0.03 | | |
| ICC | 0.05 | | |
| N ID Plasmodium | 10 | | |
| Observations | 200 | | |
| Marginal $R^2$ / Conditional $R^2$ | 0.322 / 0.353 | | |

**Table S2** Mixed-effect regression for reaction times as a function of food quality in the slime moulds study. Sclerotia identity was included as a random factor. The regression was performed using R (RStudio Version 1.2.1335; function *lmer*, package lme4). Given the typical skewness of reaction times, the dependant variable was transformed (i.e., normalized) using the *bestNormalize* function in R. As the food quality of equal alternatives increased, reaction times significantly decreased.
